# TrkB Activation During a Critical Period Mimics the Protective Effects of Early Visual Experience on Perception and the Stability of Receptive Fields in Adult Superior Colliculus

**DOI:** 10.1101/435784

**Authors:** David B. Mudd, Timothy S. Balmer, So Yeon Kim, Noura Machhour, Sarah L. Pallas

**Affiliations:** Neuroscience Institute, Georgia State University, Atlanta, GA 30303, USA

## Abstract

During a critical period in development, spontaneous and evoked retinal activity shape visual pathways in an adaptive fashion. Interestingly, spontaneous activity is sufficient for spatial refinement of visual receptive fields in superior colliculus (SC) and visual cortex (V1), but early visual experience is necessary to maintain inhibitory synapses and stabilize RFs in adulthood (Carrasco et al. 2005, 2011; Carrasco & Pallas 2006; Balmer & Pallas 2015a). In visual cortex (V1), brain-derived neurotrophic factor (BDNF) and its high affinity receptor TrkB are important for development of visual acuity, inhibition, and regulation of the critical period for ocular dominance plasticity (Hanover et al., 1999; Huang et al., 1999; Gianfranceschi et al., 2003). To examine the generality of this signaling pathway for visual system plasticity, the present study examined the role of TrkB signaling during the critical period for RF refinement in SC. Activating TrkB receptors during the critical period (P33-40) in DR subjects produced normally refined RFs, and blocking TrkB receptors in light-exposed animals resulted in enlarged adult RFs like those in DR animals. We also report here that deprivation- or TrkB blockade-induced RF enlargement in adulthood impaired fear responses to looming overhead stimuli, and negatively impacted visual acuity. Thus, early TrkB activation is both necessary and sufficient to maintain visual RF refinement, robust looming responses, and visual acuity in adulthood. These findings suggest a common signaling pathway exists for the maturation of inhibition between V1 and SC.

**Significance Statement:** Receptive field refinement in superior colliculus (SC) differs from more commonly studied examples of critical period plasticity in visual pathways in that it does not require visual experience to occur; rather spontaneous activity is sufficient. Maintenance of refinement beyond puberty requires a brief, early exposure to light in order to stabilize the lateral inhibition that shapes receptive fields. We find that TrkB activation during a critical period can substitute for visual experience in maintaining receptive field refinement into adulthood, and that this maintenance is beneficial to visual survival behaviors. Thus, as in some other types of plasticity, TrkB signaling plays a crucial role in RF refinement.

## Introduction

As sensory pathways transition from a highly plastic state early in life to a stable state in adulthood, stimulus tuning properties are progressively sharpened through neural activity-dependent plasticity and are then maintained in that state. Much of the investigation into the regulation of critical periods has centered on the development of visual system connectivity, particularly on the developmental increase of GABAergic inhibition. Development of inhibition in primary visual cortex (V1) is controlled by experience driven BDNF signaling through its high affinity receptor TrkB (Huang et al., 1999; Huang and Reichardt, 2003; Jiang et al., 2005; Gao et al., 2014), and has been shown to substitute for experience in the context of ocular dominance tuning in transgenic mice (Gianfranceschi et al., 2003). Here we address the generalizability of TrkB signaling as a mechanism underlying critical period regulation across different tuning properties and visual regions.

Receptive field (RF) refinement is an essential step in visual system development. Visual experience is required for development of acuity in monkeys (Regal et al., 1976; Teller et al., 1978), cats (Timney et al., 1978; Derrington and Hawken, 1981), and rats (Fagiolini et al., 1994), but not in V1 of mice (Prusky and Douglas, 2003; Kang et al., 2013). Our previous studies in Syrian hamsters demonstrated that early visual experience is not necessary for RF refinement in superior colliculus (SC) or V1, but is required to maintain refined adult RFs (Carrasco et al., 2005; Balmer and Pallas, 2015a). Chronic dark rearing (DR) beyond postnatal day (P) 60 results in expansion of RFs to juvenile size (Carrasco et al., 2005). Light exposure during an early postnatal critical period protects RFs against this later loss of refinement (Carrasco and Pallas, 2006; Balmer and Pallas, 2015a). Thus, in contrast to RF refinement in cats and primates (see Shatz, 1996, for review), development of refined RFs in hamster SC and V1 is independent of sensory experience, and requires vision only for maintenance. These results counter the common view that vision is required for development but not maintenance of visual receptive field properties (see Shatz, 1996, for review). They caution against over-generalization across features and species, and raise the possibility that RF refinement in hamster SC and V1 may occur through a distinct mechanism.

Because RF expansion in SC of DR adults results from a loss of inhibition (Carrasco et al., 2011; Balmer and Pallas, 2015b) we asked whether RF refinement and maintenance might occur through a mechanism other than TrkB directed inhibitory plasticity. We find, however, that TrkB activation during the critical period for RF refinement is necessary and sufficient to maintain refined RFs in SC and V1 of adults. Thus, BDNF-TrkB activity seems to be a common path through which visual experience influences the development and maturation of inhibition in the visual pathway. These findings raise the possibility that manipulating TrkB activity could reactivate plasticity in adults for therapeutic purposes, and could provide insight into the development of disorders that similarly involve the breakdown of mature connectivity stemming from an early developmental error.

## Materials and Methods

### Subjects

A total of 137 adult Syrian hamsters (*Mesocricetus auratus*) of both sexes was bred in-house and used in this study (see Table 1). Hamsters provide a valuable model for studying the developing visual system due to their robust and well-characterized visual responses and short gestation time (Pratt and Lisk, 1989). Hamsters were housed in social groups of up to 5 adults per cage in standard rodent cages, with enrichment items including nestlets and chew toys. All animals were provided ad libitum access to food and water 24 hours per day.

**Table 1.** Number of animals used throughout all experiments across treatment and control groups.

| Electrophysiology |  | Western Blotting |  | Behavior |  |
| --- | --- | --- | --- | --- | --- |
| RF size recordings | # | Pharm. treatment confirmation | # | Looming Task | # |
| Strobe reared | 3 | 7,8-DHF/DR | 9 | Normal | 9 |
| Vehicle/DR | 6 | ANA-12/Strobe | 5 | DR | 9 |
| 7,8-DHF/DR | 6 | Strobe/DR | 7 | 7,8-DHF | 6 |
| ANA-12/DR | 6 | Vehicle/DR | 11 | ANA-12 | 7 |
|  |  |  |  | Vehicle | 7 |
|  |  | Adult GAD65/GABAARa1 | # | Visual Acuity Water Maze | # |
|  |  | 7,8-DHF/DR | 7 | Normal | 10 |
|  |  | Vehicle/DR | 5 | DR | 10 |
|  |  | ANA-12/DR | 4 | 7,8-DHF | 6 |
|  |  | Normal | 6 | ANA-12 | 7 |
|  |  | DR | 4 | Vehicle | 7 |

Twenty-four adult mice (C57BL/6) of both sexes were bred in-house and used in the visual looming behavioral assay only. Mice are a valuable model for perceptual tasks because they are commonly used in behavior experiments and are widely studied animal model in visual neuroscience.

### Surgery

Electrophysiological recordings were made in sedated animals as described previously (Carrasco et al., 2005). In brief, animals were deeply anesthetized with intraperitoneal injections of urethane (2g/kg, split into 3-4 doses). Surgical levels of anesthesia were confirmed via withdrawal reflexes, respiration rate, and muscle tone, with supplemental ¼ doses of urethane given as needed. Preoperative doses of atropine (0.05 mg/kg) were administered after the onset of anesthesia to stabilize breathing and reduce secretions in the respiratory tract. A single injection of dexamethasone (1mg/kg) was used as a prophylactic anti-inflammatory. The surgical site was then shaved and cleaned with 70% ethanol, and the head was stabilized with a bite-bar restraint. A midline incision was made in the scalp to expose the skull, followed by an approximately 5mm bilateral craniotomy extending from bregma to lambda, and retraction of the meninges. The cortex and hippocampus were aspirated unilaterally to expose the underlying SC. Removal of cortex has no observable effect on SC neuron RF properties in hamsters, except for impairments in cortically-mediated direction tuning (Chalupa, 1981; Ahmadlou et al., 2017).

### Experimental design and statistical analysis

#### Light treatment groups

Normally reared hamsters or mice were housed in a 12/12 hour, reversed light-dark cycle. DR animals were housed in a darkroom, within which were several light-tight housing cabinets. Pregnant dams of DR subjects were transferred into DR housing approximately 3 days before parturition. During drug administration and for general husbandry purposes, they were briefly exposed to dim red light at a wavelength not visible to Syrian hamsters (Huhman and Albers, 1994).

To test the effect of TrkB receptor blockade on RF maintenance, strobe light exposure was used rather than a 12/12 light cycle because of the likelihood that the injected antagonist would not be effective throughout the 12-hour daily light exposure. Strobe-exposed animals were placed in a small enclosure containing a light flashing at approximately 25 Hz for 5 hours a day on each day of the critical period (7 days total). The timing of strobe exposure overlaps the 6 hour effective dose curve of the TrkB antagonist (Cazorla et al., 2011). This method exposed test subjects to a total duration of light exposure similar to the amount that was sufficient to maintain RF refinement in SC (Carrasco et al., 2005).

#### Drug treatment groups

In order to test the hypothesis that TrkB activation is necessary and/or sufficient for adult maintenance of RF refinement, all animals were administered a daily injection of either a TrkB agonist (Andero et al., 2011), a TrkB antagonist (Mui et al., 2018), or vehicle during the critical period for RF maturation in SC (P33-40) under dim red light conditions (Carrasco et al., 2005). Specifically, treatment consisted of an intraperitoneal injection of either the TrkB agonist 7,8-Dihydroxyflavone (7,8-DHF) (98% Sigma Aldrich CAS#: 38183-03-8) (10mg/kg) or the antagonist ANA-12 (Sigma Aldrich #SML0209) 0.15g/ml, made fresh daily, dissolved in diluted dimethylsulfoxide (60% DMSO in DI water), or DMSO alone as a negative control. Animals were weighed prior to each daily injection to ensure that the 1 mg/kg dose of drug or vehicle remained consistent throughout the treatment phase.

#### Statistical analysis

A Student’s *t*-test or a One-Way Analysis of Variance (ANOVA), followed by post-hoc Bonferroni tests were used to compare parametric data with equal variance between groups and a normally distributed data set. Descriptive statistics for these analyses are provided as mean ± standard error of the mean (SEM). For data not meeting these criteria, a Mann-Whitney rank sum test or a Kruskal-Wallis One-way ANOVA on ranks was used, followed by a Dunn’s post hoc test, with data presented as median ± interquartile range (IQR).

### Western blotting

Animals were euthanized with a sodium pentobarbital-and phenytoin sodium-containing mixture (Euthasol >150 mg/kg IP). Brains were immediately extracted and flash frozen in cold 2-methylbutane on dry ice, then stored at −80°C or immediately dissected for preparation of lysates. Individual left and right tecta were excised and lysed in RIPA buffer (150mM NaCl, 150mM Tris, 1% NP-40, 0.1% SDS, 0.5% sodium deoxycholate) containing 2% Halt protease inhibitor (ThermoFisher Scientific). Proteins were visualized using SuperSignal West Pico Chemiluminescent Substrate kits (Life Technologies) and imaged on an ImageQuant LAS4000 mini imaging system (GE Healthcare Life Sciences), or IRdye fluorescent secondaries (Li-Cor), imaged on an Odyssey CLx fluorescent imaging system (Li-Cor). Protein levels were quantified as the optical density of the phosphorylated TrkB proteins relative to the optical density of total TrkB protein using ImageJ. No difference was detected between the two imaging methods using identical membranes, thus the data were combined. To assess the effectiveness of the TrkB agonist and antagonist, 33 animals received the drug doses IP, and then either remained in their DR habitat or were exposed to strobe conditions for 2 hours, followed by euthanasia and tissue harvest. Rabbit anti-pTrkB (Y817) (1:1000, Abcam Cat # ab81288), and rabbit anti-pan (total) TrkB (80G2, 1:500, Cell Signaling Technologies Cat# 4607) were used to confirm that the drugs were having the expected effect on TrkB phosphorylation in *in vivo*. The Y817 phosphorylation site was chosen because it is a BDNF phosphorylation site (Liu et al., 2014) and its phosphorylation is a reliable marker for calcium release (Hubbard and Miller, 2007). Phosphorylation of Y817 also activates protein kinase C (PKC), which is associated with activity dependent synaptogenesis in visual cortical development (Zhang et al., 2005). Negative controls included lanes with primary antibody but no protein to confirm specificity of the bands identified at the targeted molecular weight.

### Assessment of pre- and post-synaptic inhibitory signaling strength

To characterize and compare treatment dependent changes in inhibitory signaling in adulthood we examined levels of GAD-65 and GABB_A_R α1 protein using antibodies (mouse anti-GAD-65 1:10 (Developmental Studies Hybridoma Bank-University of Iowa GAD-6), and rabbit anti-GABAARα1 1:1000 (Abcam ab33299)). Negative controls included lanes without protein and lanes without primary antibody.

### Electrophysiology

Single unit extracellular recordings were obtained with Teflon coated, glass insulated microelectrodes (Kation Scientific, Catalog: W1011-7, 2-2.3 MΩ). The electrode was positioned perpendicular to the exposed SC, and lowered into the tissue using a Kopf Model 650 micropositioner. All recordings were obtained within 200 µm of the surface of the SC to ensure they were made from the superficial, retinorecipient layers. Electrical signals were amplified, filtered (10,000x; 0.5-3kHz; Bak Electronics A-1), and digitized at 20 kHz using Spike2 software and CED hardware (Micro 1401-2; Cambridge Electronic Design).

### Visual stimulus presentation

In order to measure RF size, we used a visual stimulus presented on a CRT monitor (60Hz refresh rate) positioned 40 cm from the left eye. The monitor was maintained at its highest contrast and brightness settings for each experiment. Visual stimulus generation was accomplished using custom MATLAB (Mathworks) software with the PsychToolBox-3 application. The visual stimulus consisted of a bright white square traveling from dorsal to ventral visual field at 14°/s across a black background. The stimulus size was 1 degree in diameter and each vertical traverse shifted 2° along the x axis of the monitor after each presentation, with a 3 second inter-stimulus interval, as in (Balmer and Pallas, 2015a).

### Analysis of RFs

Spike2 software (Cambridge Electronic Design) was used for offline spike sorting of single units (approximately four unique visually responsive neurons were isolated per recording site). Analysis of RF size was completed by a researcher blind to treatment group. Receptive field diameter along the azimuthal axis was measured by plotting the visual field location from which spiking responses were produced as the stimulus vertically traversed the monitor, from nasal to temporal visual field. A uniform fraction of the peak response (20%) was defined as the minimum stimulus-evoked response threshold, as in a previous study (Balmer and Pallas, 2015a). Responses were normalized by setting the peak response of each single unit to 1.0 to account for differences in response strength between individual units. RF size data were compared between treatment groups to quantify the effect of TrkB manipulation vs. vehicle treatment. Data for LR Control and DR Control groups were taken from our previously published study, using the same methods (Carrasco et al., 2005).

### Looming response task

An open-top box with dimensions of 58.7cm long × 42.9cm wide and 32.4cm in height was used to test the fear reflex to an expanding spot approaching from above. The test subjects were light and dark reared Syrian hamsters and C57BL/6J mice, aged >P90, housed either in 12h:12h light:dark cycle or in 24h dark. Five trials were conducted under white light, in the animals’ subjective daytime between the hours of 1900-2200. Both groups were exposed to white light for less than 90 minutes per trial. Alcohol (70%) was used to clean the apparatus before and between each trial, to eliminate olfactory cues. A plastic cup was spray painted black and offered as a hiding chamber. Each subject was placed in the center of the apparatus with the white monitor screen above and given at least 10 minutes for acclimation. The visual stimuli were programmed and displayed using the Psychtoolbox module for MATLAB. A spherical, black, looming stimulus on a white background was initiated once the subject was out of the hiding chamber and in the center of the arena. The stimulus expanded from 3.5 degrees of visual angle to 56.5 degrees in 2.35 seconds, remained at that size for 250ms, and then restarted the sequence with a 250ms delay. The fear response was considered positive if the subject either froze or fled into the cup within 5 seconds of the stimulus initiation. The fear response was considered negative if the subject did not demonstrate any freezing or fleeing behavior.

### Visual water maze task

A two-alternative, forced-choice visual discrimination task was used to assess the spatial acuity of Syrian hamsters across all treatment groups. The task consisted of a trapezoidal shaped Y maze half-submerged in a pool with 15cm of tepid (22°C) water. A hidden escape platform that was submerged in one of the two distal arms of the maze was separated by an opaque divider 40cm in length, with the far end of the divider marking the decision line. Identical monitors (Dell Model 1707FPt) placed side by side and above the distal ends of the maze displayed either a gray screen or a sinusoidal grating of vertical black and white bars. The maximum detectable number of grating cycles (cycles per degree – cpd) that occurred throughout the span of a single visual degree for the subjects was calculated and used as a measure of visual acuity. Screen reflections on the surface of the water hid the platform when viewed from water level. Hamsters were trained to escape from the Y maze by swimming toward the screen displaying the gratings, where the hidden platform was located. Visual acuity was determined by increasing the number of grating cycles across the screen (adding 1 complete cycle for each progressive set of trials). The escape platform location was randomly determined before each set of trials. When an incorrect choice was made, the animal would be assayed over several more trials to determine an overall success rate at that cpd. If the animal fell below a 70% success rate then its preliminary visual acuity was determined to be the total cpd of the previous trial set. The final visual acuity value for each animal was progressively narrowed down over the course of several days and approximately 60 trials per animal.

### Data sharing information

Intellectual property rights are set by Georgia State University Policy No. GSU: 4.00.08. Data will be embargoed only until publication, unless the University requests a delay in public dissemination if necessary to permit the University to secure protection for Intellectual Property disclosed to it by the PI. After publication, we are willing to share any of the data used to generate our manuscripts as long as the PI and members of the Pallas lab involved in generating the data and the funding sources receive proper attribution.

## Results

The developmental transition from plastic to stable receptive field properties maintains the activity driven changes that occur in early life. Early visual experience is required for refined RFs to be stabilized and thus maintained into adulthood (Carrasco et al., 2005), but it remains unclear what molecular changes are responsible. BDNF protects against degradation of visual acuity in visual cortex of dark reared mice (Gianfranceschi et al., 2003), and we examined whether BDNF-TrkB signaling might also be protective of acuity in SC of dark reared hamsters, in which case it may be a general mechanism through which sensory experience has its maturational effects across species and brain area. Alternatively, adult maintenance of RF refinement in Syrian hamsters may occur through a different signaling pathway.

### 7,8-DHF and ANA-12 are both effective modulators of TrkB receptors throughout the visual midbrain

In the study on the effects of visual deprivation on visual cortical development of mice mentioned above (Gianfranceschi et al., 2003), a genetic approach was used for constitutive over-expression of BDNF. We asked a more time-limited question- whether increasing signaling through the TrkB BDNF receptor only during an early critical period would have a similar effect. To accomplish this, we used a pharmacological approach that allowed us to control the timing of TrkB manipulation. We reasoned that if increased TrkB signaling that is limited to the critical period could rescue receptive field properties from the effects of visual deprivation, it would suggest that a common mechanism exists for stabilizing inhibitory synapses across different visual brain areas and species, despite the differences in timing. Systemic injection of the isoflavone 7,8-DHF as a TrkB receptor agonist (Andero et al., 2011; Liu et al., 2014), and ANA-12 as an antagonist (Lawson et al., 2014; Ren et al., 2015) have been used to manipulate TrkB activation in previous studies. In order to determine whether we could use these drugs to achieve a level of TrkB activation during the critical period that would be comparable to that provided by light exposure, we assayed their ability to phosphorylate and dephosphorylate TrkB receptors at the same site that is phosphorylated by light exposure and BDNF binding (Y817) (Poo, 2001; Hubbard and Miller, 2007; Liu et al., 2014). Test subjects from each treatment group (7,8-DHF + DR, n=9; Strobe + ANA-12, n=5; Strobe alone, n=7; Vehicle + DR, n=11) were euthanized 3 hours after receiving treatment, and the brains were collected for processing.. We then used Western blotting to measure the amount of activated (pTrkB) relative to total TrkB from V1, SC, and hippocampus (as a non-retinorecipient control region). We found that the pharmacological manipulations intended to stimulate TrkB receptors were working as intended, in that immunoblotting with antibodies against phosphorylated (activated) and total TrkB receptors revealed strong, treatment-induced increases in TrkB phosphorylation at Y817 throughout the brain (**Figure 2)**. In all three areas 7,8-DHF had a robust effect on increasing TrkB phosphorylation in DR subjects well beyond that of the Vehicle + DR injection group; SC (F(3,24) = 12.503, p < 0.001, ANOVA) V1 (F(3,12) = 24.757 p <0.001, ANOVA), hippocampus (F(3,12) = 6.070 p = 0.009, ANOVA). Visual experience (strobe) also induced increases in pTrkB in both visual brain areas compared to vehicle: SC (0.523 ± 0.0916, p = 0.046, n=5), V1 (0.529 ± 0.0812, p = 0.004, n=4), but not in hippocampus, as expected for a non-visual area. Conversely, treatment with the TrkB antagonist ANA-12 + strobe reduced pTrkB levels in relation to total TrkB in both SC (0.132±0.243, p = 0.034 n=3) and V1 (0.098±0.0265 p<0.001 n=4). These findings support our claim that pharmacological activation of TrkB agonists can modulate TrkB activity in hamster SC, similar to short, stroboscopic light treatments, whereas antagonist treatment can reduce TrkB activation, mimicking visual deprivation.

**Figure 1.**
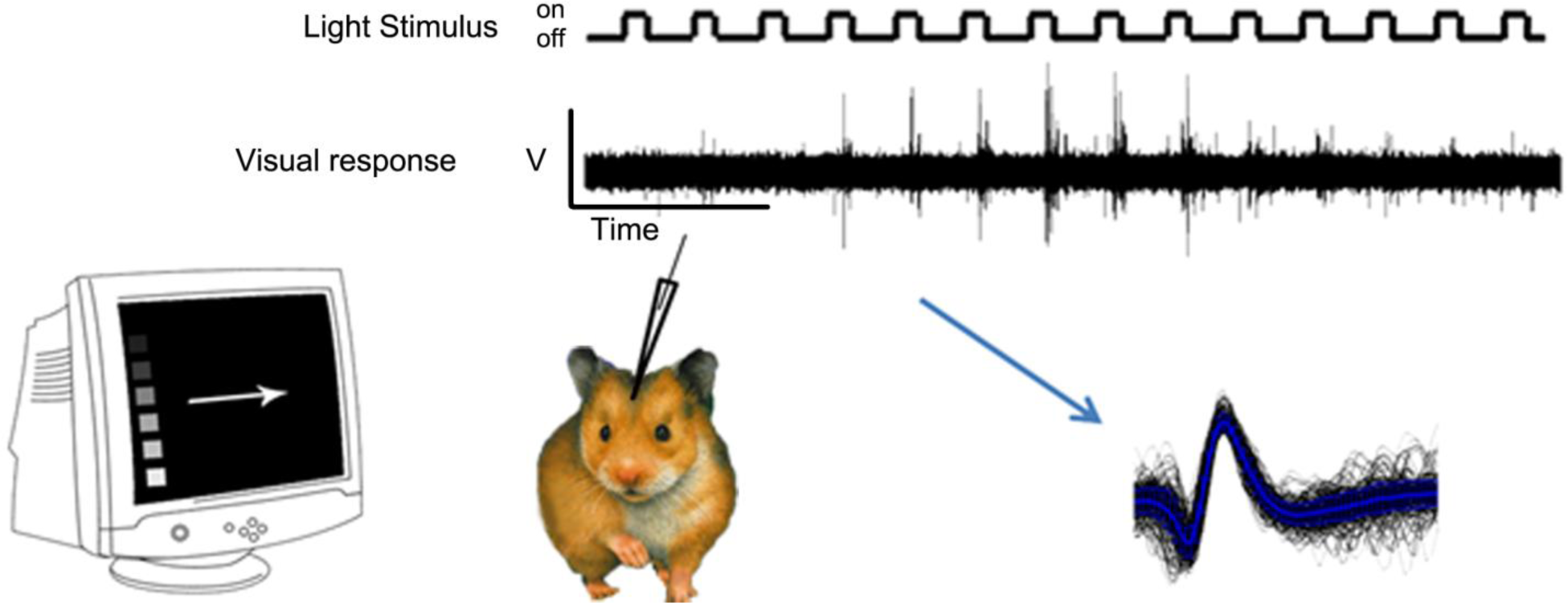
Graphical description of experimental procedure for measuring visual RF sizes with *in vivo*, extracellular, single unit recordings of stimulus-evoked action potentials in SC.

**Figure 2.**
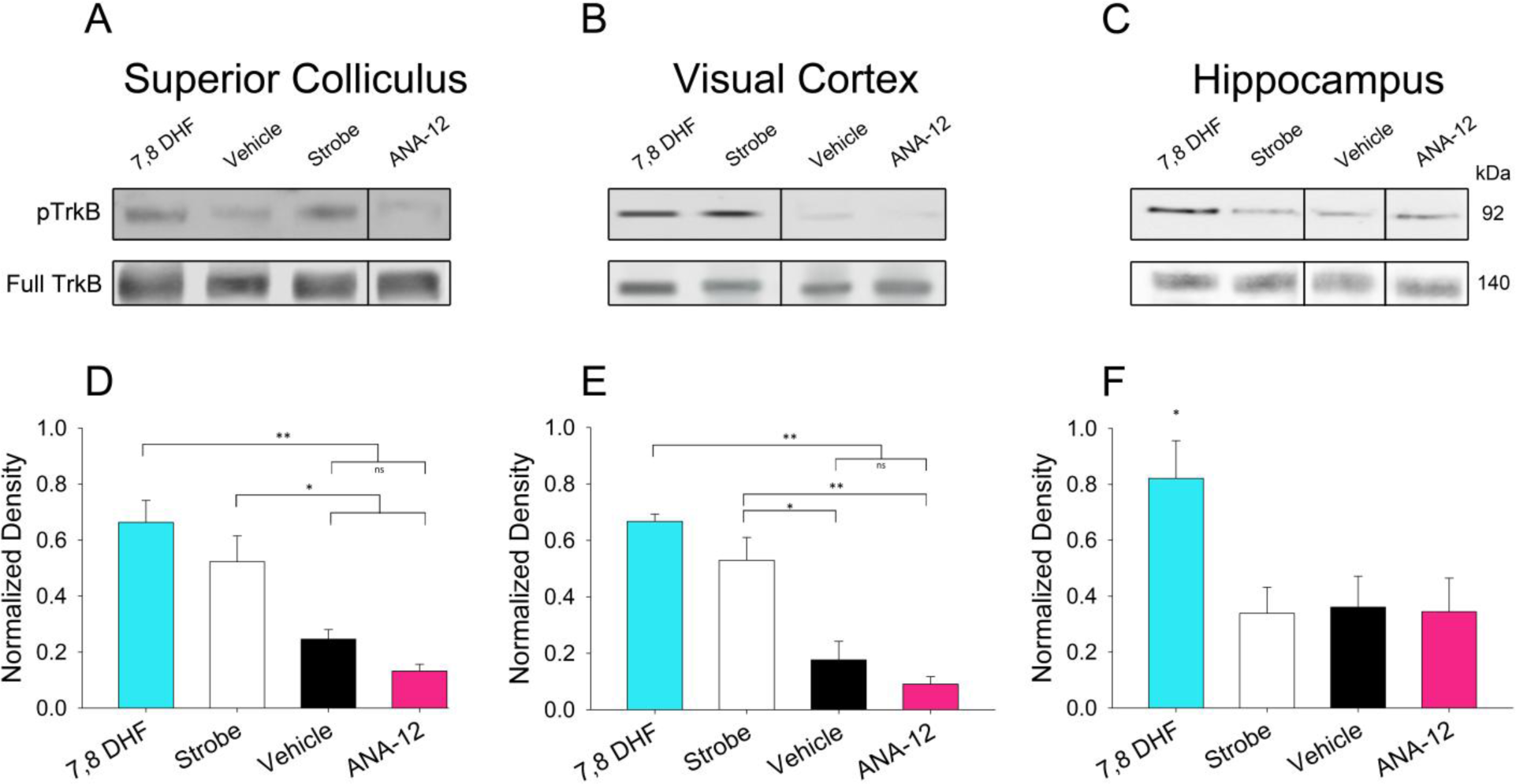
Drug treatments were effective at modulating TrkB activity in SC, V1, and Hippocampus. 7,8-DHF administration from P33-40 increased TrkB receptor activation, and ANA-12 treatment blocked TrkB activation, in all brain areas assayed. **(A-C)** Example blots of treatment groups generated using 20 µg of protein per lane. All presented lanes are from the same gels, with nonadjacent lanes revealed by vertical lines between them. **(D-F)** Densitometric analyses of Western blots generated from SC **(D)** V1 **(E)** or Hippocampus **(F)** lysates prepared from juvenile hamsters (∼P33) receiving the TrkB receptor agonist (7,8-DHF), visual experience (strobe light for 1 hour), strobe light + the TrkB antagonist ANA-12, or vehicle injection revealed differences in activation levels of TrkB receptors between groups. Agonist and strobe light exposure greatly increased levels of phosphorylated (p)TrkB in SC compared to vehicle. Analysis of ANA-12 treatment on TrkB phosphorylation during one hour of strobe light exposure revealed that the antagonist is effective in preventing visual experience-evoked TrkB activation. The density of the anti- pTrkB (Y817) band is normalized to the anti-total TrkB (80G2) protein band to measure differences in TrkB activity. Data presented as mean ± SEM. *p<0.05, **p<0.01.

### Elevating TrkB receptor phosphorylation levels during the critical period maintains SC receptive field refinement into adulthood

In SC and V1, refinement of RFs during postnatal development occurs independently of visual experience, but maintaining refined RF size into adulthood requires visual experience during an early critical period (Carrasco et al., 2005; Balmer and Pallas, 2015a). Because genetically increasing BDNF expression throughout life rescues RF size in dark reared visual cortex of mice (Gianfranceschi et al., 2003), we reasoned that early BDNF signaling might also be involved in RF maintenance in adult superior colliculus. To investigate whether TrkB activation during the critical period for RF plasticity has the same effect on RF size as visual experience, we pharmacologically manipulated TrkB activation during the critical period and measured RF sizes in adult superior colliculus (>P90). Data for LR Control and DR Control groups were taken from our previously published study, using the same methods (Carrasco et al., 2005). Our approach was to dark rear hamsters from <P0 to >P90 and provide daily treatment with the TrkB agonist 7,8-DHF throughout the critical period (P33-P40). As predicted, DR hamsters that were treated with the agonist maintained a significantly smaller RF size (12° ± 6°, n = 92) compared to vehicle injected control animals (18° ± 4°, p<0.05, n = 84) (H(3) = 120.118 p = <0.001, Kruskal Wallis One-Way ANOVA on Ranks) **(Figure 3)**. Importantly, these are measurements of single unit RF sizes, and not overlapping, adjacent RFs that might be measured from multiunit extracellular recordings. These results support the hypothesis that TrkB activation during the critical period is sufficient to maintain RF refinement in SC into adulthood.

**Figure 3.**
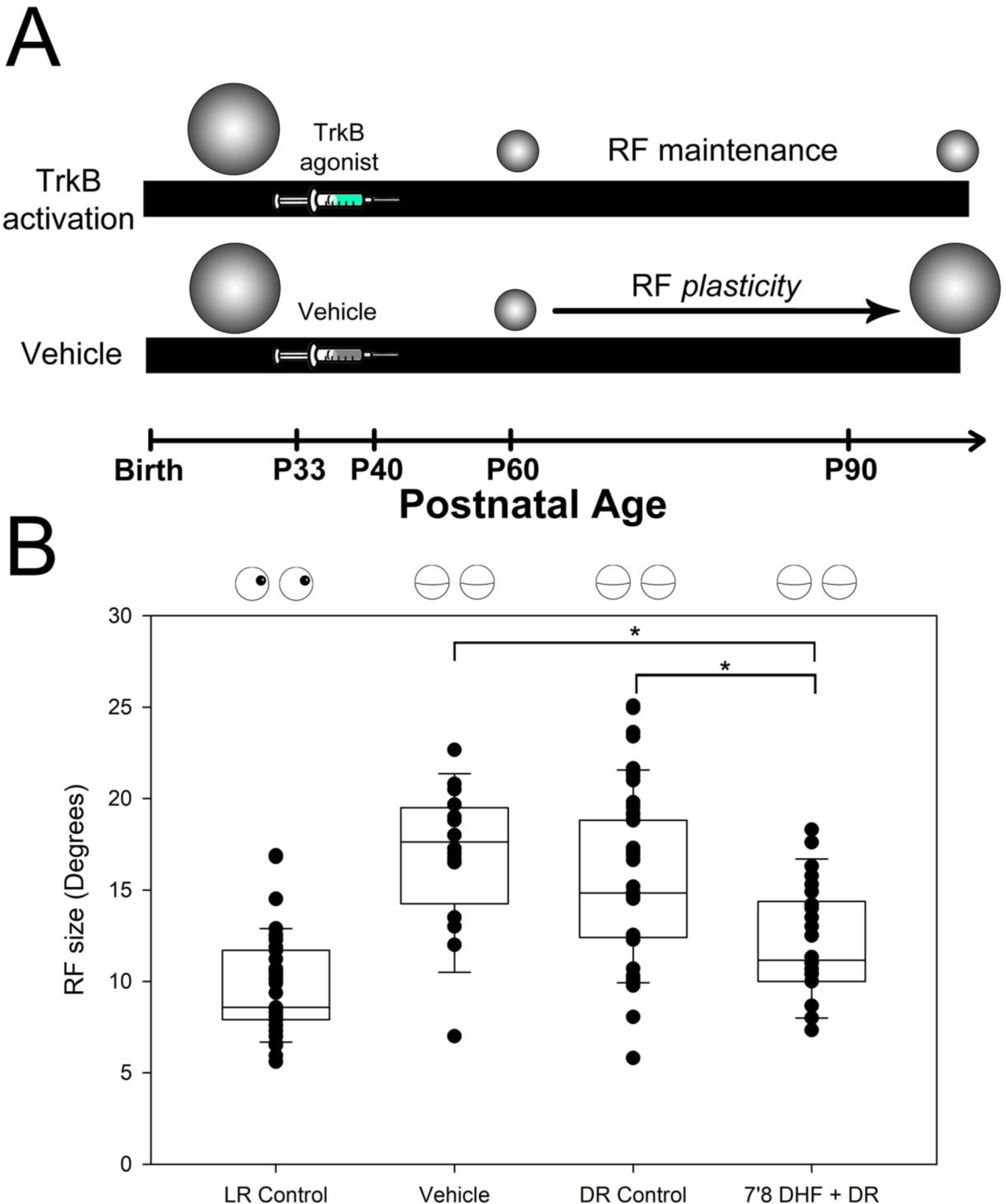
TrkB activation during the critical period maintains RF refinement into late (>P90) adulthood. **(A)** Experimental design and summary of findings. **(B)** RF sizes for each experimental group measured in visual degrees and plotted as individual data points. LR and DR Control data are from Carrasco et al. (2005). Open pair of eyes across the top of graph indicates the group was given visual experience. Closed eyes indicate group was dark reared throughout development. Data presented as median ± IQR. *p<0.05.

### Decreasing TrkB phosphorylation levels during the critical period prevents maintenance of SC receptive field refinement into adulthood

Next, we tested whether TrkB activation is a requirement for maintenance of refined RFs. Our approach was to use the TrkB antagonist ANA-12 to block the light-induced phosphorylation of TrkB receptors that occurs during visual experience. ANA-12 was administered during a stroboscopic presentation of light for 5 hr/day throughout the critical period (P33-P40), followed by return to the dark room until adulthood (>P90). This stroboscopic light treatment was sufficient to maintain RF refinement in SC into adulthood **(Figure 4).** Single-unit recordings from SC neurons in animals receiving the antagonist revealed significantly larger RFs (20° ± 4° (n=82 neurons)) than in neurons from strobe light treated animals (12° ± 4° (n=29 neurons)) **(Figure 4B)** and interestingly, larger RFs than in vehicle injected, light exposed subjects (18° ± 4° n = 84)(H(3) = 153.50, p = < 0.001, Kruskal Wallis One-Way ANOVA on Ranks). We further compared the effects of all treatments on adult RF size across different quadrants of the SC and observed no significant differences between strobe, vehicle + DR, or ANA-12 + strobe, suggesting that no regions of the SC are particularly susceptible to DR. Note that the plot includes data from LR Control and DR Control groups in our previously published study (Carrasco et al., 2005). Interestingly, we did observe more refinement in RFs located in the rostral (10° ± 2.5°, n= 21) and medial (10° ± 4°, n=17) quadrants of the SC compared to the caudal (14° ± 4°, n=29) and lateral (14° ± 4°, n= 28) quadrants **(Figure 5)**, consistent with recent findings that certain visual stimulus features may be sampled more robustly at different regions of the visual field (El-Danaf and Huberman, 2019). These findings further supports the hypothesis that TrkB activity is responsible for maintenance of RF refinement, because blocking TrkB activity during the critical period resulted in enlarged RFs in adulthood, despite adequate visual experience for maintenance of refined receptive fields.

**Figure 4.**
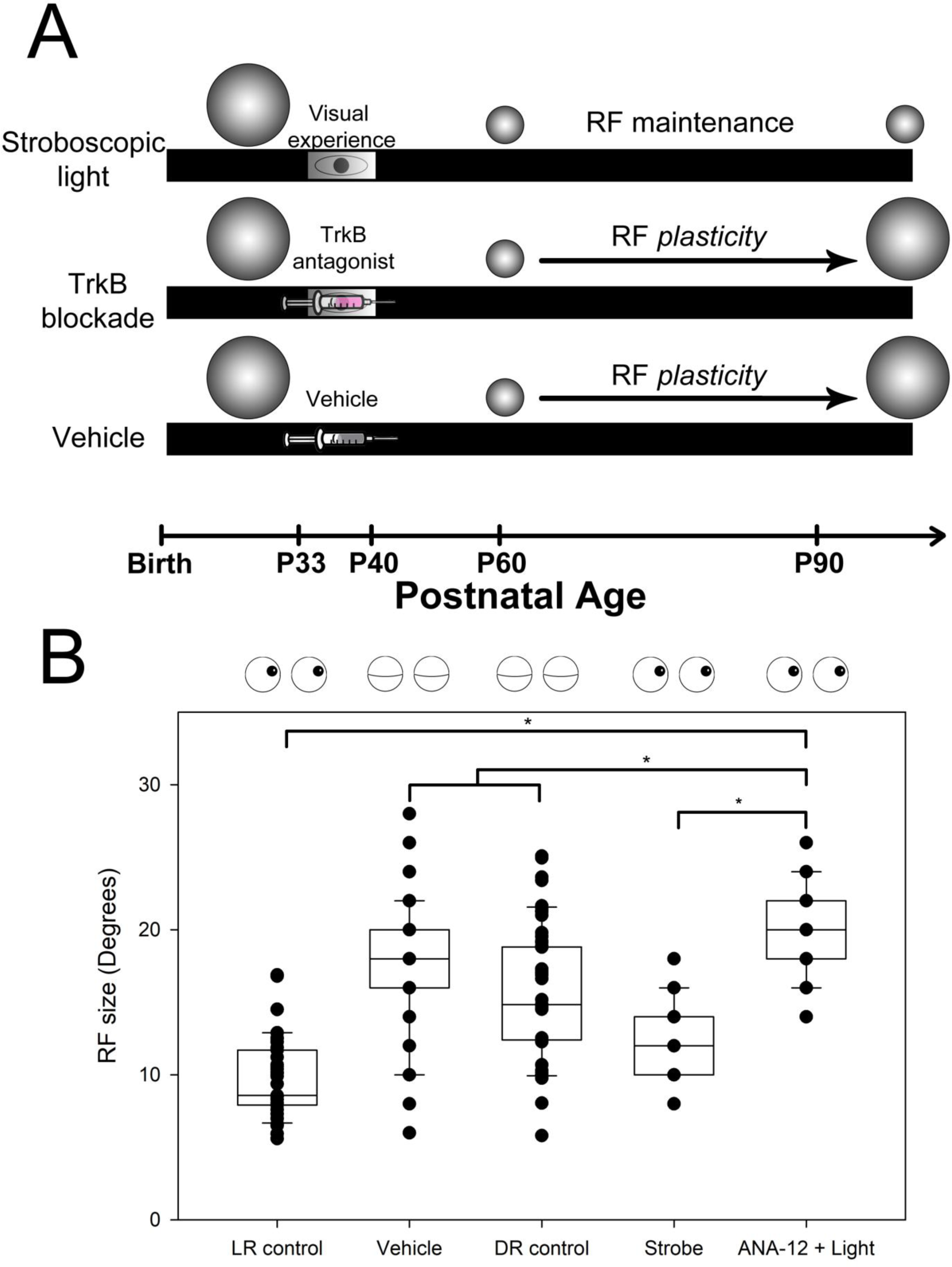
TrkB blockade during the critical period blocks the protective effects of visual experience for SC RF refinement in adulthood (>P90). DR animals were given sufficient stroboscopic visual experience during the critical period to maintain RF refinement in SC, but TrkB blockade during that time period blocked the protective effects of light. **(A)** Experimental timeline for treatment and resulting RF changes during development. **(B)** RF sizes for each experimental group measured in visual degrees and plotted as individual data points. LR and DR Control data are from Carrasco et al. (2005). Symbols as in Figure 3. Data presented as median ± IQR. *p < 0.05.

**Figure 5.**
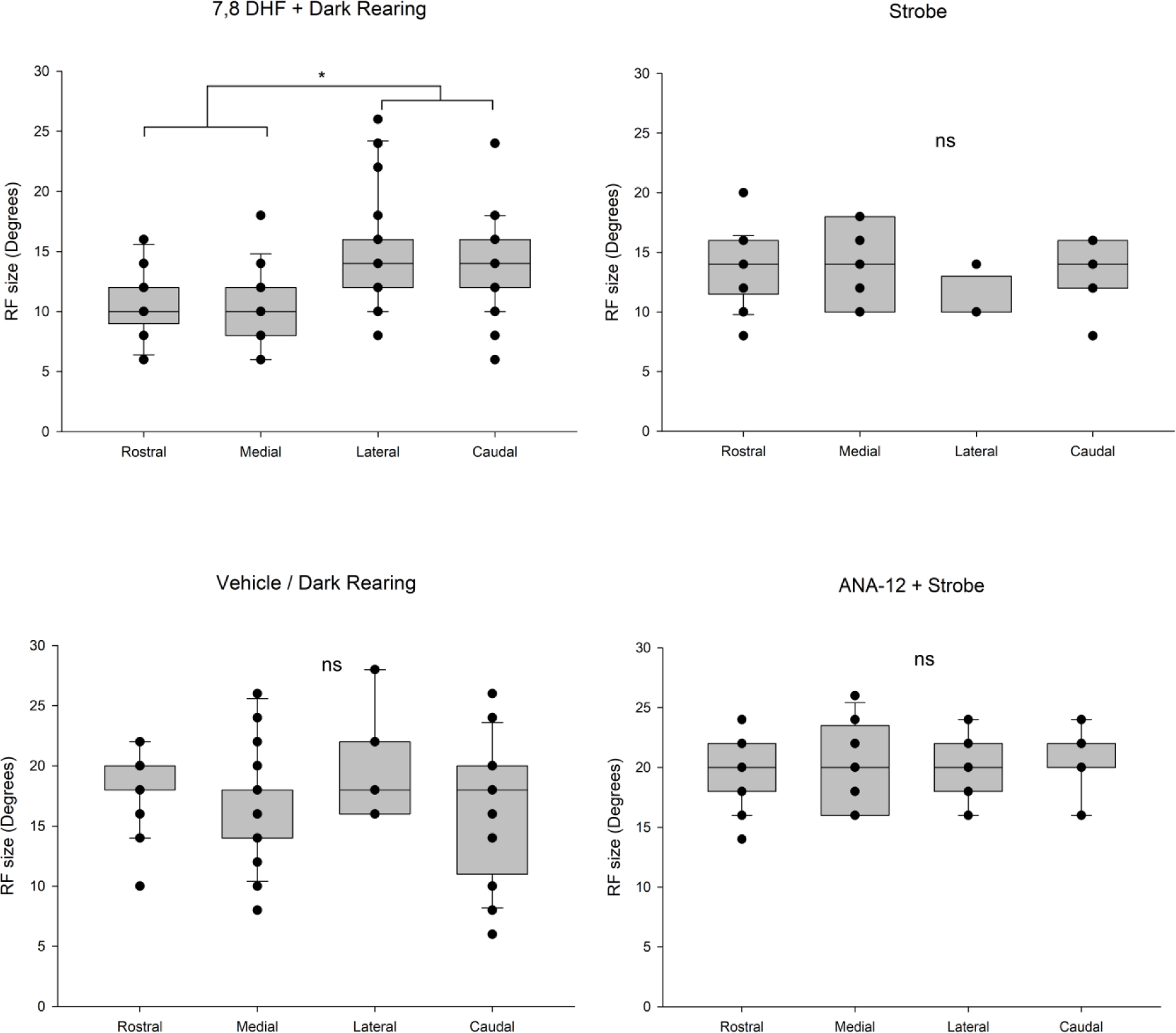
Analysis of treatment effect on RF size across quadrants of SC. In the 7,8-DHF groups the RFs in SC neurons representing the rostral and medial quadrants of the visual field were smaller than those in the caudal and lateral quadrants. In adult hamsters, these regions receive topographic input from the nasal and dorsal visual fields, respectively. No other treatment elicited a quadrant specific effect on RF size. Data presented as median ± IQR. *p< 0.05.

### TrkB activity during the critical period preserves RF refinement by maintaining adult levels of inhibition

Results from our previous investigations suggested that loss of inhibition could account for the RF enlargement in adult DR animals (Carrasco et al., 2011; Balmer and Pallas, 2015a). If TrkB is the pathway through which visual deprivation leads to a loss of lateral inhibition in SC and thus enlarged RFs in adulthood, then reduced TrkB activation should result in reduced inhibition. Thus we examined how altering TrkB activation affected GABA and GABAA receptor levels. Examinations of the GABA precursor enzyme (GAD65) levels and GABA_A_R receptor expression levels in TrkB agonist and antagonist-treated hamsters were used to examine any potential changes in adult inhibition. Assays of GAD65 and GABAA receptor levels allowed us to address if increasing TrkB activity during the critical period in DR hamsters would maintain adult levels of lateral inhibition in SC in the same manner as visual experience, or if TrkB activity is affecting RF refinement in SC through a different mechanism.

We measured adult (>P90) levels of GAD65 and GABA_A_Rα1 receptor subunit protein in drug-treated and control animals using Western blotting. Agonist (7,8-DHF/DR) treated animals had higher relative cytoplasmic GAD65 expression (0.216 ± 0.006, n = 7 animals) compared to vehicle /DR treated (0.104 ± 0.0121, n = 5), and antagonist (ANA-12/strobe) treated hamsters (0.085 ± 0.0092, n=4) (F(4,28) = 102.747, p < 0.001, One-Way ANOVA). Normally reared and DR control groups performed as expected, with normally reared hamsters having much higher GAD65 expression compared to DR hamsters (Normal: 0.242 ± 0.0057, n = 6; DR: 0.0737 ± 0.0048, n=4; One-Way ANOVA, p<0.001) **(Figure 6A)**. In contrast, drug treatment had no effect on GABA_A_Rα1 expression (p = 0.97) **(Figure 6B)**. Presynaptic GAD65 expression was reduced in groups in which RFs had become unrefined (Vehicle, DR, ANA-12/Strobe), whereas post-synaptic receptor function was not affected. These results suggest that visual experience and TrkB activity are not functioning via unique signaling pathways.

**Figure 6.**
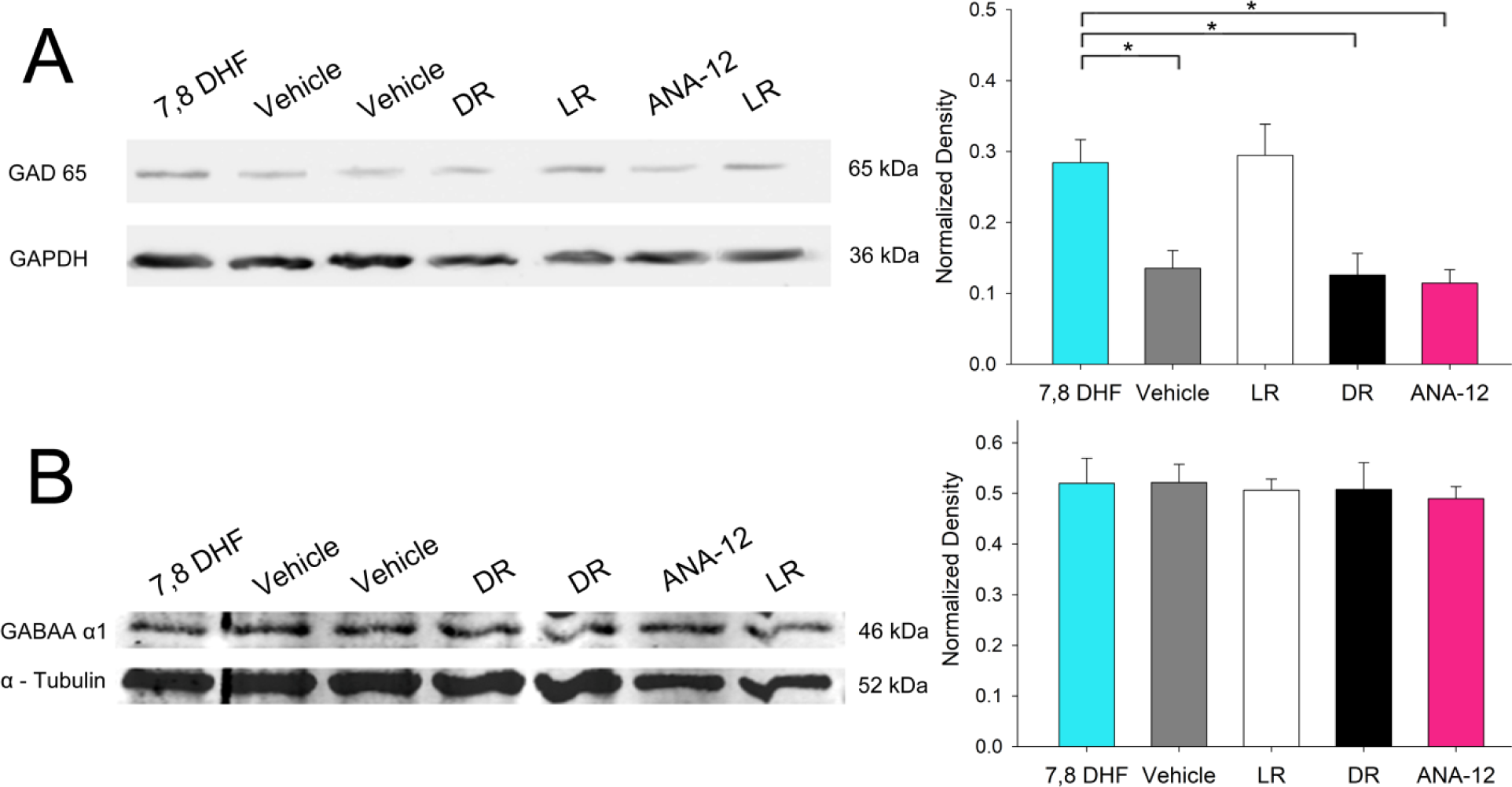
Early TrkB expression is both necessary and sufficient to maintain increased presynaptic inhibition in adult SC. **(A)** Adult GAD65 expression levels were maintained in TrkB agonist injected animals but decreased in DR, vehicle, and ANA-12 injected animals. **(B)** GABA_A_R α1 expression remained constant across all treatment groups and rearing conditions. Data presented as mean relative optical density ± SEM. *p<0.001.

### Dark rearing impairs fear responses to looming visual stimuli

RF refinement is critical for the development of visual acuity and environmental awareness. DR animals have a number of visual deficits in cortex such as poorer visual acuity and broader orientation and direction tuning (Fagiolini et al., 1994), but deficits in SC have rarely been characterized. In rodents, the retino-SC pathway is arguably more relevant to visual behavior than the geniculocortical pathway (Sherman and Spear, 1982; Li et al., 2015; Beltramo and Scanziani, 2019), especially compared to predators. We hypothesized that refined RFs are necessary for SC dependent visual behaviors to function in adulthood, and predicted that groups with enlarged RF’s (DR/Vehicle, ANA-12/Strobe) would have impaired task performance. We tested this hypothesis in adult hamsters by examining differences in fear responses (escape or freezing behavior; see Methods) to visual looming stimuli **(Figure 7A, B)**, an SC dependent behavior (Zhao et al., 2014; Shang et al., 2018) that is dependent on input from retinal W3 cells (Zhang et al., 2012). We found that DR (40% ± 6%, n=8), vehicle (23% ± 8%, n=7), and ANA-12/strobe (26% ± 4%, n=7) treated hamsters were less likely to respond to overhead looming stimuli than normally reared (78% ± 6%, n=8), or 7,8-DHF treated (70% ± 4%, n=6) hamsters **(Figure 7C, E)** (F(4, 35) = 17.73, p<0.001, One-way ANOVA).. Dark rearing/larger RFs had a particularly robust effect on the escape (“flight” to shelter) behavior, with very few occurrences of flight in groups in which RFs have expanded in adulthood **(Figure 7C)**. These data suggest that the failure to maintain RF refinement in adult SC has a negative impact on instinctual fear responses to a looming visual object.

**Figure 7.**
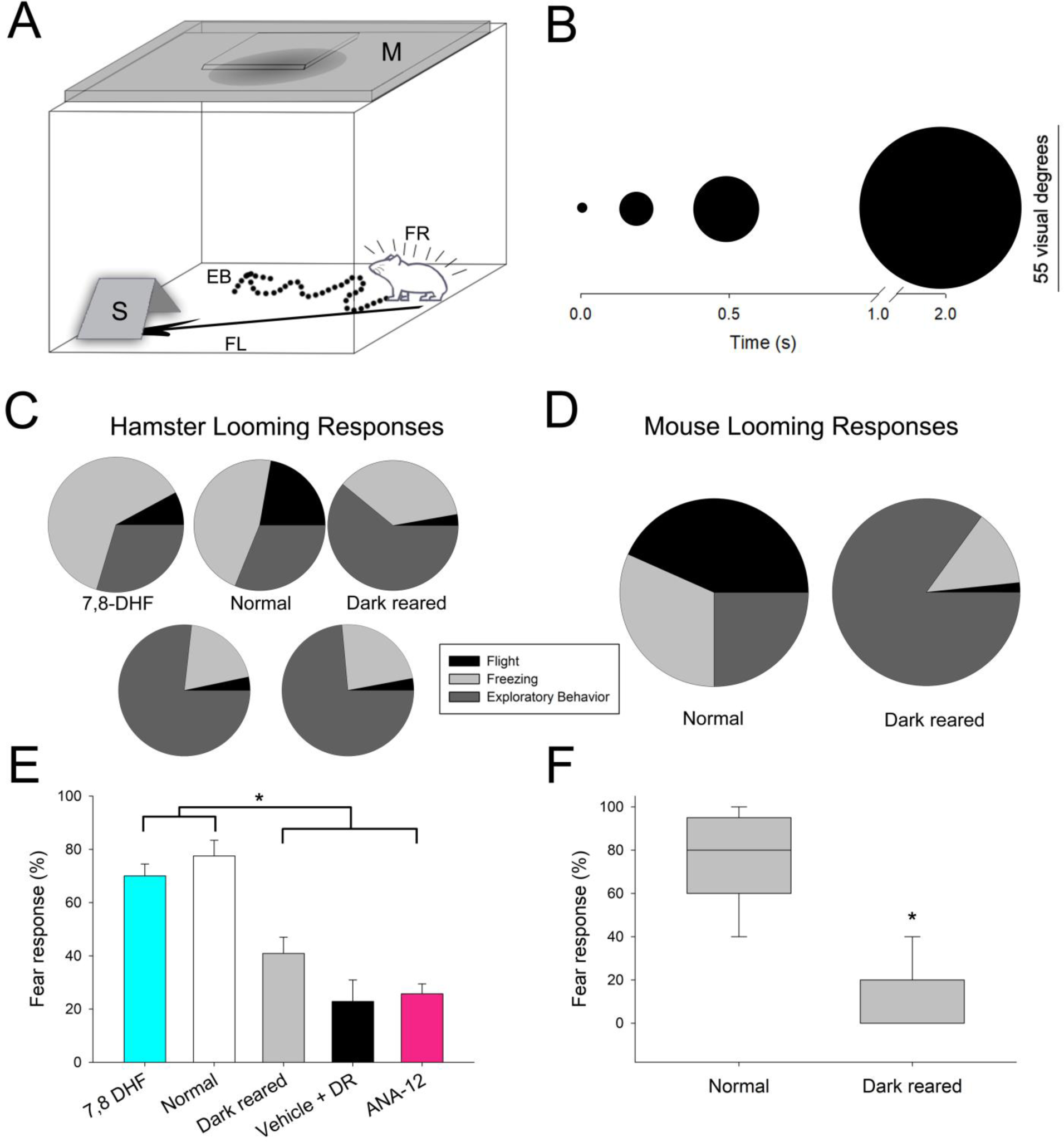
Dark rearing and subsequent enlargement of RF size in SC reduces fear responses to looming stimuli in hamsters and mice. **(A)** Schematic of apparatus. A box with a monitor (M) suspended above it projecting the looming stimulus, and a shelter (S) placed at the far end. Animals responded to looming stimuli either by fleeing (FL) into the shelter or freezing (FR) in place. Unresponsive animals continued exploratory behavior (EB). **(B)** Expansion of the looming stimulus from the start of a cycle to the end (approximately 2s). **(C/D)** Occurrences of each response type to looming stimuli per animal in each experiment group for 5s after stimulus presentation in hamsters (C) and mice (D). Responses were determined on an ascending scale from **exploratory behavior (EB)** < **freezing** < **flight**, with only the highest observed behavior reported for each trial. **(E/F)** The frequency of an escape response (freezing/flight) to looming stimuli compared between normally reared and DR groups in hamsters (E) and mice (F). Data presented as mean fear response ± SEM (E) and median ± IQR (F). *p <0.05.

It is important to address the possibility of species-specific responses to light deprivation. Because hamsters are a crepuscular species (Apfelbach and Wester, 1977) and as such encounter their environment in both light and darkness, the effects of light deprivation may not be as detrimental as on a nocturnal species that is less reliant on vision for survival. To test this hypothesis, we carried out the same visual perceptual test on mice that had been normally reared or DR. We found that the effect of DR on the frequency of fear responses to looming stimuli was similar between mice and hamsters. As with DR hamsters, DR mice were less likely to respond to overhead looming stimuli (20%± 20%, n=12) than normally reared mice (80% ± 30%, n=12) (T=220, n(small)=12, n(big)=12, p<0.001 Mann-Whitney Rank Sum Test,).. Surprisingly, the decrease in fear responses in DR mice was even greater than the decrease in DR hamsters **(Figure 7D, F)**, contrary to what might be predicted in a nocturnal species like mice. These data suggest that both hamsters and mice are susceptible to DR induced disruptions of visual perception in SC. They also support our contention that RF size refinement is an important event in visual development and could have a detrimental effect on behaviors that are important for survival.

### TrkB activity during the critical period for RF refinement is both necessary and sufficient for visual acuity improvements to persist into adulthood

In rats, cortex-dependent visual acuity at eye opening, as assessed by visual evoked potentials, is less than half that in adulthood (Fagiolini et al., 1994). In both SC and V1 of hamsters, chronic DR impairs the stability of the refined RFs, resulting in RF expansion in adulthood (Carrasco et al., 2005; Balmer and Pallas, 2015a) We hypothesized that TrkB activity during the critical period for RF refinement would be both necessary and sufficient for adult visual acuity to be preserved. To test this hypothesis, we compared the proficiency of hamsters in performing a spatial discrimination task in adulthood across treatment groups (7,8-DHF/DR, ANA-12/Strobe, Strobe, DR, and Vehicle/DR) **(Figure 8A,B)**. We found that visual acuity was similar between strobe treated (0.695 cpd ±0.01, n=9), and 7,8-DHF/DR treated (0.704 cpd ± 0.01, n=6) hamsters, but was reduced in Vehicle/DR (0.473 cpd ± 0.01, n=7), DR (0.481 ± 0.01, n=11), and ANA-12/Strobe treated hamsters (0.516 ± 0.01, n=7, F(4,35) = 42.31, p < 0.001, One-Way ANOVA) **(Figure 8C,D)**. These results demonstrate that TrkB activity during the critical period for RF refinement is both necessary and sufficient for high visual acuity in adulthood, and that RF size refinement is important for overall visual acuity and for survival behaviors such as the looming response.

**Figure 8.**
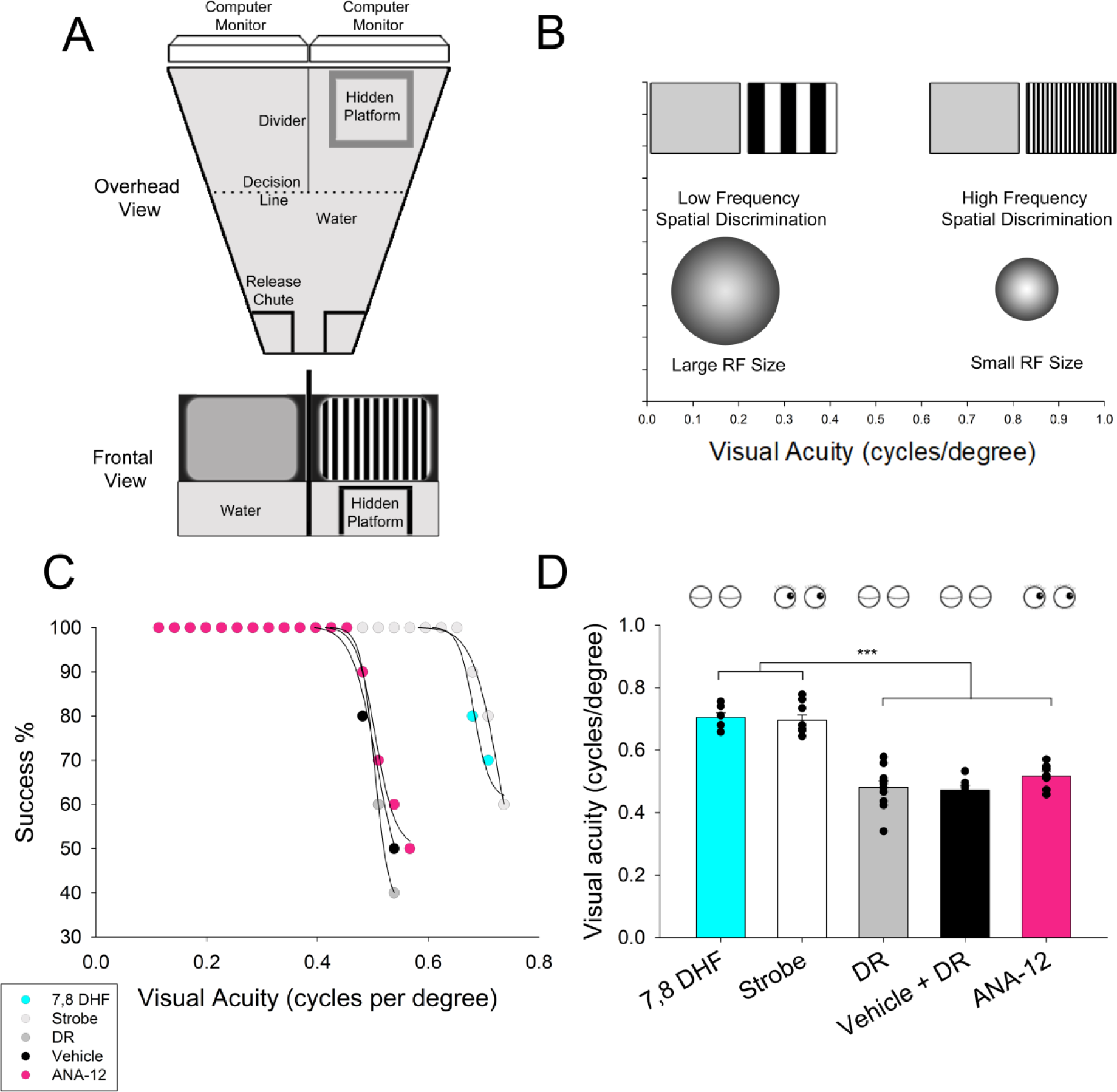
TrkB activity during the critical period for RF refinement maintains adult levels of visual acuity. **(A)** Schematic of apparatus from two vantage points. A trapezoidal Y maze filled with 15cm of water was used in a forced choice behavioral assay. Monitors placed above each arm displayed either a vertically oriented sine-wave grating to indicate the location of the escape platform or a gray screen, in random order. **(B)** Progression of trials testing visual discrimination against increasing frequencies of sine-wave gratings. Animals with larger RFs are expected to have poorer visual discrimination ability. **(C)** Acuity performance curves were applied to each treatment group. Each point represents the average success rate at the indicated spatial frequency. Acuity was determined by identifying the point where the curve crossed the horizontal line indicating a 70% success rate. (D) Comparison of visual acuities in cpd across all treatment groups. Symbols as in Figure 3. Data presented as mean ± SEM. ***p < 0.001.

Together, these results demonstrate that TrkB activation can substitute for visual experience during the critical period of RF plasticity, and they support the hypothesis that TrkB activation is both necessary and sufficient for the maintenance of RF refinement in SC **(Fig. 9)**. Our results also provide evidence that this early increase in TrkB expression reduces presynaptic GAD65 expression in adult SC. In addition, RF refinement in SC is important for visual behavior, in that larger RFs result in impaired responsivity to looming visual stimuli, and poorer discrimination of spatial gratings.

**Figure 9.**
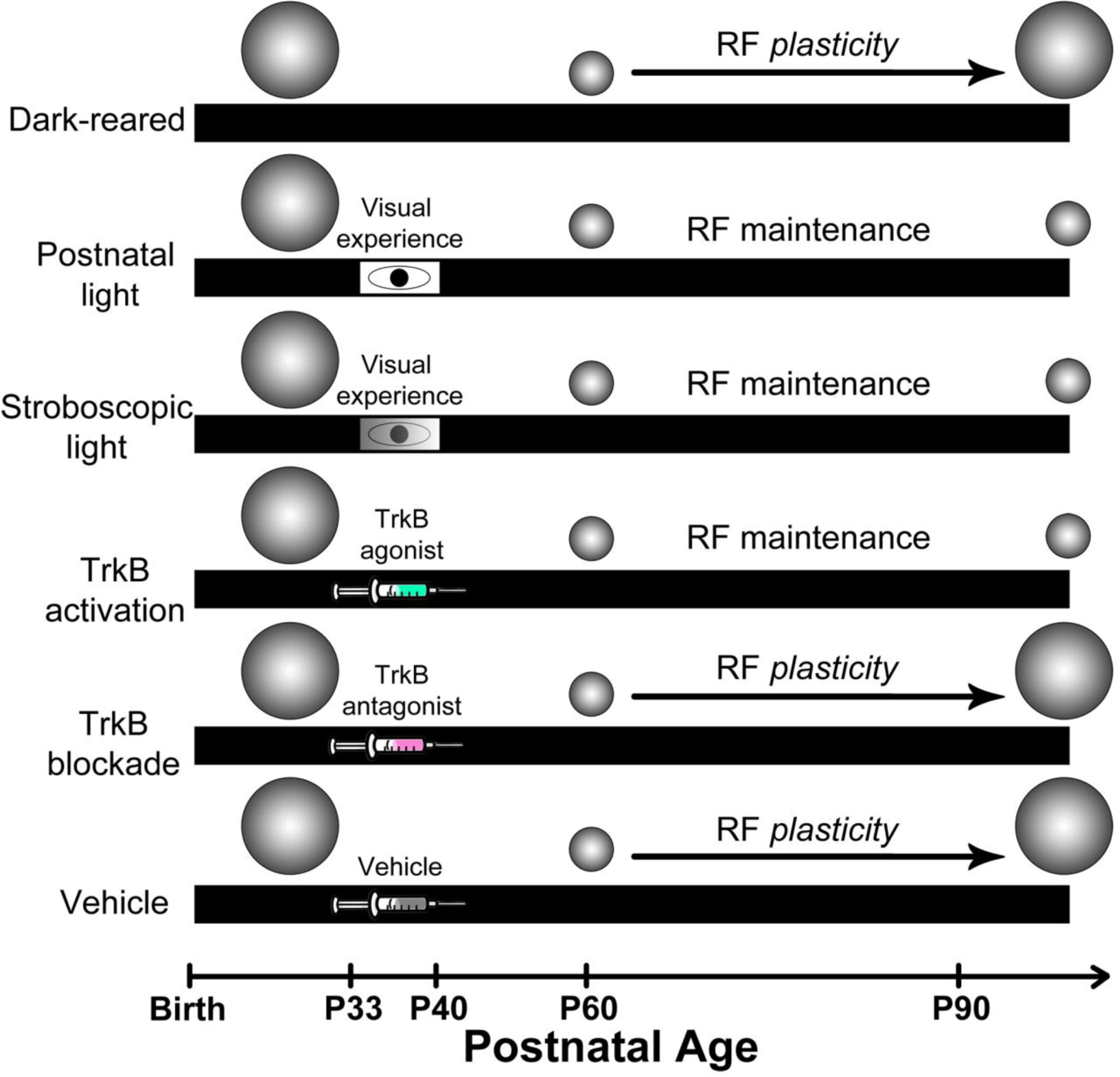
Summary. Cartoon showing findings regarding the dependence of RF size maintenance in adult SC on early visual experience, and the role that TrkB activation plays within that process.

## Discussion

Previously we reported that spontaneous activity alone is sufficient for RF refinement in SC and V1 (Carrasco and Pallas, 2006; Balmer and Pallas, 2015a). However, light exposure for several days during an early critical period is necessary for maintaining refinement into adulthood (Carrasco et al., 2005). These results were surprising for two reasons; first because vision is thought to be necessary for development but not for adult maintenance of function, and secondly because early deprivation did not have any detectable effect until adulthood. These previous results provided the rationale for the current study of the mechanism through which early visual experience maintains RF size past puberty and into adulthood.

Visual deprivation can permanently impair the development of some stimulus tuning properties, yet have little effect on others, depending on species. For example, orientation selectivity in visual cortex will begin to appear in juvenile DR ferrets (Chapman and Stryker, 1993; Chapman et al., 1996; Chalupa and Snider, 1998; Issa et al., 1999; White et al., 2001) but fails to sharpen to mature levels (Huberman et al., 2008) and direction selectivity in V1 fails to develop at all (Li et al., 2006; Van Hooser et al., 2012). Additionally, ocular dominance columns in V1 form in DR cats (Wiesel and Hubel, 1974; Horton and Hocking, 1996), but ocular dominance plasticity is prolonged into adulthood as a result (Mower et al., 1981; Cynader, 1983). In contrast, some RF properties develop without visual experience in mice (Rochefort et al., 2011), but continued deprivation degrades tuning in adult mice and rats (Hensch, 2005). This suggests that one role of early visual experience may be to fine tune and then and stabilize previously established connections in their mature state. This maturation is often associated with increased GABAergic inhibition (Hensch et al., 1998; Fagiolini et al., 2004) initiated by increases in visually evoked BDNF and subsequent TrkB receptor activation (Huang et al., 1999). Recent work further connects early visual experience, BDNF activity, and cortical maturation persisting into adulthood (Zhang et al., 2018), though BDNF was reported to have an inverse relationship with early cortical maturation during the critical period (P23-28). We report that increasing TrkB phosphorylation during a critical period can substitute for visual experience and forestall the negative effects of visual deprivation on RF refinement. These findings clarify the requisite role of experience-driven TrkB activity in stabilizing adult RFs and preventing deleterious adult plasticity. They suggest that TrkB receptor signaling is the convergence point through which visual activity drives the maturation of inhibition in the superior colliculus and visual cortex, at least in hamsters and mice (Gianfranceschi et al., 2003; Balmer and Pallas, 2015a).

### TrkB signaling mediates activity-dependent maturation of visual processing circuits in V1 and SC

Development of ocular dominance columns is subject to a critical period during which monocular deprivation can shift the representation of cortical cells away from the closed eye (Wiesel and Hubel, 1963, 1965). Dark rearing causes a prolongation of the critical period for ocular dominance plasticity, perhaps as a result of prolonged immaturity of NMDA receptors (Carmignoto and Vicini, 1992) and GABAergic neurons (Jiang et al., 2005). In contrast, RF refinement in SC and V1 of hamsters and rats is unaffected by visual deprivation, and early experience is necessary only for stabilizing RFs in adulthood. Visual experience is thought to influence ocular dominance development through a BDNF-mediated signaling pathway that promotes maturation of inhibitory synapses and regulation of critical period plasticity. Visual input drives NMDA receptor activity, increasing BDNF levels and thus triggering maturation of inhibition (Castren et al., 1992; Greenberg et al., 2009; Park and Poo, 2013). Transcription factors such as Npas4 regulate genes associated with plasticity, including BDNF (Lin et al., 2008; Van Hooser et al., 2012; Bloodgood et al., 2013). BDNF in turn binds to and activates TrkB receptors (Pollock et al., 2001; Viegi et al., 2002), promoting the expression of GAD, GABA, and GABAA receptors throughout the brain (Rutherford et al., 1997; Huang et al., 1999; Jovanovic et al., 2004; Porcher et al., 2011; Sanchez-Huertas and Rico, 2011). Critical period closure and the ensuing restriction of visual cortical plasticity are associated with visual experience-induced changes in NMDA and GABA receptor composition (Carmignoto and Vicini, 1992; Fox et al., 1992; Stocca and Vicini, 1998; Chen et al., 2001; Li et al., 2017), increases in GABA expression (Jiang et al., 2005), and perineuronal net development (Sur et al., 1988; Bavelier et al., 2010; Beurdeley et al., 2012; Ye and Miao, 2013; Wen et al., 2018). Spontaneous activity is not sufficient to drive these changes in the context of ocular dominance plasticity (Sur et al., 1988; Huberman et al., 2008; Chalupa, 2009).

The similar effects of TrkB signaling on RF refinement in both SC and V1 are surprising for a number of reasons. V1 requires several more (P33-P40) visual experience to stabilize adult RF refinement (>P90) than SC (P37-P40) (Balmer and Pallas, 2015a). V1 also has different GABAergic cell classes than SC, and very few of the parvalbumin positive interneurons (Mize, 1992; Choi et al., 2009; Villalobos et al., 2018) that are essential in V1 ocular dominance plasticity (Hensch, 2005). Thus, it seems likely that there are some undiscovered differences in the TrkB signaling pathway downstream of TrkB activation.

### How does early TrkB signaling maintain RF refinement?

Visual experience-regulated TrkB activity may function in either a permissive or an instructive role at different stages of visual system development. In the permissive role scenario, (Huang et al., 1999; Seki et al., 2003) experience driven activity increases overall BDNF production and subsequent TrkB activity. The TrkB activity would then permit maturation of the GABAergic synapses necessary to develop or stabilize RF properties. In the instructive role scenario (Kossel et al., 2001), visual experience would drive TrkB activity to form specific ensembles of visual neurons that respond to different types of visual input. Our finding that early TrkB activity maintains RF refinement in adult SC suggests that the BDNF-TrkB signaling pathway plays more of a permissive than an instructive role. This view is supported by our previous result that RF refinement will occur without any visual experience in SC (Carrasco et al., 2005) and in V1 (Balmer and Pallas, 2015a) and that the loss of refinement in DR adults coincides with reductions in GABA expression and postsynaptic GABA receptor function. Thus RF refinement does not require instructive signaling from the BDNF-TrkB signaling pathway to occur, but rather requires permissive signaling to maintain its stability long term.

### How does early visual experience contribute to survival?

In early life, most mammals have underdeveloped sensory and motor capabilities, and require intensive parental care to survive. As they age, sensory and motor abilities mature in response to intrinsic maturational processes as well as in response to environmental stimuli, leading to improved overall function and independent survival in adulthood. Vision is one of the most important senses for many mammals, because it facilitates object and feature detection for purposes of conspecific and interspecific interactions, as well as for feeding and locomotion (Thinus-Blanc, 1996). Improvements in visual acuity, as well as orientation, size, and motion tuning, are all important for identifying objects in the environment, and require varying amounts of early visual experience to develop (see Huberman et al., 2008, for review). In SC, the multisensory integration of vision with other senses considerably enhances the salience of a sensory event (Meredith and Stein, 1983; Frens and Van Opstal, 1998) and is important for orientation behaviors (Stein et al., 1989). Among these behaviors, looming stimulus detection and fear responses allow the avoidance of aerial predators (Yilmaz and Meister, 2013). Although the function of looming detection is well understood, the contribution of TrkB activity to development of this behavior during early development has yet to be explored. Our results show how early visual experience, increased TrkB signaling, and refined RFs in SC are essential for the performance of looming stimulus detection and the defensive behaviors associated with it. Surprisingly, our results indicate that looming response behaviors in mice, a nocturnal species with presumably less reliance on vision, were more detrimentally affected by dark rearing compared to hamsters. This could be due to poorer overall visual acuity, or perhaps reduced dorsal visual field coverage in mice, something we will explore in future work. Interestingly, of the two different fear response behaviors assessed here, the freezing response was reduced, and the escape response was nearly eliminated in DR animals. Freezing is uniquely effective as a defense from aerial predators (Eilam, 2005; De Franceschi et al., 2016), and our finding that it becomes the dominant response for DR animals raises the possibility that it could be a more instinctual behavior than escape responses. Another possibility is that dark rearing alters the downstream processing pathway that facilitates the behavioral response. For example, the SC-ventral midline thalamus connection processes the overall reaction to visual threats (Salay et al., 2018), and could be affected by downstream TrkB activity. Future experiments will address these unanswered questions and further our understanding of TrkB signaling during early development.

## Acknowledgements

Support for this work was provided by a GSU Brains and Behavior Fellowship and a GSU Center for Neuromics student grant awarded to D.B.M., and a GSU Brains and Behavior Seed grant, a National Science Foundation grant (IOS-1656838) and a DARPA grant (HR0011-18-2-0019, TA2) awarded to S.L.P. We would like to thank Dr. Angela Mabb of the GSU Neuroscience department and Dr. Hyuk-Kyu Seoh of the GSU Biology department for providing guidance on Western Blot imaging and analysis, members of the Pallas lab for technical support and manuscript review, and the Department of Animal Resources staff at GSU for excellent animal care.

